# Large Scale Functional and Effective Connectivity Alterations cross the Huntington’s Disease Integrated Staging System

**DOI:** 10.64898/2025.11.30.691364

**Authors:** Murat Demirtas, Dorian Pustina, Andrew Wood, Cristina Sampaio, Jakub Vohryzek, Gustavo Deco

**Affiliations:** Center for Brain and Cognition, Universitat Pompeu Fabra, Barcelona; CHDI Management, Inc., the company that manages the scientific activities of CHDI Foundation, Inc., Princeton, NJ, USA

**Keywords:** Huntington’s Disease, Huntington’s Disease Integrated Staging System, TRACK-ON HD, Resting-state fMRI, Functional connectivity, Effective connectivity, Computational Modeling

## Abstract

Huntington’s disease (HD) is a progressive neurodegenerative disease with severe motor, cognitive and behavioral symptoms. There is a recent impetus to develop treatments that slow progression before clinical signs emerge. Such early interventions require biomarkers sensitive to the very earliest HD progression. Here we applied cutting-edge fMRI analysis on data collected in the Track-On HD study to evaluate whether functional and effective connectivity obtained from model-free and model-based approaches can produce useful biomarkers of HD progression. We analyzed data from 231 participants with up to three annual visits each and created five groups comprising normative controls and four HD groups according to the Huntington’s Disease Integrated Staging System (HD-ISS). We found significant differences across the HD-ISS stages but not in their longitudinal change. Specifically, we found attenuated functional and effective connectivity in the caudate nucleus in HD-ISS-2, and this effect extended to other corticostriatal connections in HD-ISS-3. Overall, most of the alterations were only evident in advanced HD stages, and we did not observe widespread alterations in cortical connectivity and graph topography. Although HD-ISS groups did not differ in the amount of in-scanner motion, we did find measures of functional and effective connectivity to be sensitive to motion. We conclude that fMRI can indeed capture attenuation of cortico-striatal functional and effective connectivity across HD progression. This is the first study to investigate fMRI alterations across the recently created HD-ISS stages within a novel framework of signal flow across the whole brain.

## Introduction

Huntington’s disease (HD) is a debilitating progressive neurodegenerative disease transmitted through monogenic autosomal-dominant inheritance of a pathologically-expanded polyglutamine-coding trinucleotide (CAG) repeat in exon 1 of the huntingtin (*HTT*) gene (Macdonald, 1993). Signs/symptoms encompass a range of motor, cognitive, and behavioral abnormalities, typically emerging between 40 and 50 years of age. The age of sign/symptom emergence is dependent upon the CAG-repeat length. Nevertheless, this emergence involves substantial individual variation. The neuropathological mechanisms of the disease are not fully understood; however, there is clear evidence that the striatal structures—specifically the caudate and putamen—are critically affected by the disease (Jiang et al., 2023; Tabrizi et al., 2009). Medium spiny neurons in the striatum are lost at a higher rate than other types of neurons, and this degeneration is reflected in the macroscopic volumetric atrophy measured with MRI (Kinnunen et al., 2021). Caudate and putamen atrophy is detected many years before the emergence of clinical signs (Scahill et al., 2025) and has recently been selected as the only biological marker occurring before clinical motor onset in the Huntington’s Disease Integrated Staging System (HD-ISS) (Tabrizi et al., 2022). There are currently no approved disease-modifying therapeutics for HD, which causes death within 10-20 years from the initial onset of symptoms.

The HD-ISS has expanded the traditional view of HD progression by including early events before the emergence of clinical signs and symptoms. Specifically, Stage 0 indicates individuals who carry the pathologic *HTT* gene, Stage 1 indicates biological degeneration reflected in atrophy of caudate or putamen, Stage 2 indicates the presence of clinical signs in motor or cognitive domains, and Stage 3 indicates loss of everyday functional abilities. This complete characterization of the disease, from birth to loss of function, has opened up the possibility of developing therapeutics that prevent degeneration rather than reversing pathological processes. Early intervention has a better chance of halting or slowing the progression of the disease, but this will require early biomarkers around Stages 0 and 1 that can measure significant change when there are no clinical signs and no measurable clinical benefit.

Numerous publications have found anomalies in cortical and subcortical resting-state functional connectivity (rs-FC) in people with HD (PwHD) (for a review, see (Pini et al., 2020). Subcortical areas, caudate and putamen in particular, often show the most prominent impairment, not surprising given the structural degeneration of these structures (Paulsen et al., 2004). Decreased connectivity in the bilateral putamen is reported to be correlated with CAG-repeat length and with the severity of motor dysfunction (i.e., the Unified Huntington’s Disease Rating Scale, UHDRS) after clinical motor diagnosis (CMD), and it can also predict the onset of future clinical signs in early HD (Unschuld et al., 2012; Wolf et al., 2008). Another approach of functional magnetic resonance imaging (fMRI) studies is to investigate the activity of large networks composed of multiple regions that activate and deactivate together. Various studies have reported alterations in these resting-state networks (RSNs) during HD progression (Dumas et al., 2013; Werner et al., 2014). Among those alterations, the early ones are observed in the sensory-motor network (SMN) before CMD, and the later ones are found in visual and attention networks. Previous studies also reported significant topological changes in network organization (Gargouri et al., 2016; Harrington et al., 2015). In an exhaustive literature review, Pini et al. (2020) noted that the resting-state functional connectivity alterations follow a non-linear course based on the cortical topography: in somatomotor areas, PwHD exhibit hypo-connectivity before CMD, but this becomes hyper-connectivity in later-stage individuals with signs and symptoms after CMD. In frontal brain areas responsible for executive functions, the pattern is unclear, as both hypo- and hyper-connectivity have been reported in PwHD after CMD.

Most of the previous HD studies have focused on purely data-driven connectivity analyses between brain regions or networks. There has been no attempt to use large-scale modeling of the entire brain to investigate the complex interactions among multiple brain regions and their roles in the network. Such large-scale interactions can shed light on the mechanisms of HD progression and, at the same time, enable the monitoring of disease progression through specific markers (Deco and Kringelbach, 2014). Generative effective connectivity (GEC) is a framework that infers the underlying causal connectivity patterns for the observed whole-brain functional connectivity (Deco et al., 2024, 2023; Kringelbach et al., 2023). The GEC framework has been successfully implemented in Alzheimer’s disease and under naturalistic-experimental conditions (Deco et al., 2024; Demirtaş et al., 2019, 2017; Kringelbach et al., 2023). GEC has two main advantages: it filters out features of the functional connectivity that the proposed connectivity model cannot explain, and it provides a directed connectivity structure to estimate the effect of region A on region B, which may detect subtle anomalies in the directional flow of information between brain regions.

In this study we sought to explore the functional connectivity alterations using GEC and graph theory measures to test their biomarker power. We utilized data from the well-known Track-On HD study, and, given the increasing adoption of HD-ISS by the research community, we employ this staging system to classify participants into progression groups.

## Material and Methods

### Study population

We utilized data from the Track-On HD study (Tabrizi et al., 2011, 2009), which was harmonized in BIDS format and preprocessed with fMRIprep (Pustina et al., 2024). Data were available for 240 participants with a maximum of three visits each. The time gap between the first visit and the second visit was 362,97 (STD: 67.13) days, while the gap between the first and the third visits was 709.36 (STD: 58.77) days. Of the 240 participants, 128 were PwHD, and 112 of them were normative controls. In the following paragraphs, we use the terms “visit” and “session” interchangeably.

Following recommendations in the literature, we removed scans with framewise displacement larger than 0.5 (Power et al., 2012). Upon inspection of the final timeseries, we observed evident artifactual spikes in some subjects. To investigate this further, we examined the sessions in which the signal derivative was of unknown origin and exceeded 6 standard deviations. If these deviations were observed in the first or last 10 volumes, we excluded those volumes from the analysis and retained the session. However, if these deviations occurred during the recordings, it was not possible to remove them due to the short recording time of the data. Therefore, we excluded these individuals from the analyses. Following this procedure, 235 individuals remained with at least one rs-fMRI session.

No HD-ISS classification could be made for 4 individuals; therefore these participants were excluded from the analyses. The remaining 231 individuals were classified as 108 HCs, 21 HD-ISS-0, 13 HD-ISS-1, 60 HD-ISS-2 and 28 HD-ISS-3 (Table 1).

**Table 1.** Demographics of the individuals contributing to the study. The study comprised 3 rs-fMRI sessions; Baseline, 12 months and 24 months. PwHD were classified into 4 HD-ISS categories: HD-ISS-0, HD-ISS-1, HD-ISS-2, HD-ISS-3. The top of each row shows the ANOVA statistics for each variable.

| TIME OF VISIT | ISS | N | AGE | SEX<br>(% FEMALE) | FRAMEWISE<br>DISPLACEMENT | SDMT | TFC | TMS | CAG | CAP<br>SCORE |
| --- | --- | --- | --- | --- | --- | --- | --- | --- | --- | --- |
| BASELINE |  | 213 | F = 3.754,<br>p = 0.0057 | F = 2.137,<br>P = 0.077 | F = 0.569,<br>p = 0.6857 | F = 9.9192,<br>p < 0.0001 | F = 134,<br>p < 0.0001 | F = 70,<br>p < 0.0001 | F = 2.88,<br>p < 0.040 | F = 6.27,<br>p = 0.0005 |
|  | HC | 101 | 47,62 | 60,40% | 0,18 | 57,92 | 13,00 | 1,39 |  |  |
|  | HD-ISS-0 | 29 | 42,16 | 65,52% | 0,17 | 58,38 | 13,00 | 3,41 | 42,52 | 79,45 |
|  | HD-ISS-1 | 28 | 45,78 | 35,71% | 0,18 | 54,07 | 13,00 | 4,11 | 42,71 | 87,29 |
|  | HD-ISS-2 | 38 | 41,29 | 44,74% | 0,16 | 49,24 | 13,00 | 9,95 | 43,97 | 85,99 |
|  | HD-ISS-3 | 17 | 45,35 | 58,82% | 0,20 | 45,88 | 11,59 | 14,53 | 43,29 | 91,55 |
| 12 MONTHS |  | 205 | F = 6.494,<br>p < 0.0001 | F = 3.797,<br>P = 0.0053 | F = 1.700,<br>p = 0.1514 | F = 8.849,<br>p < 0.0001 | F = 43,<br>p < 0.0001 | F = 77,<br>p < 0.0001 | F = 0.405,<br>p = 0.75 | F = 7.29,<br>p = 0.0001 |
|  | HC | 96 | 49,71 | 62,50% | 0,19 | 57,53 | 12,96 | 1,58 |  |  |
|  | HD-ISS-0 | 27 | 40,70 | 77,78% | 0,16 | 61,15 | 12,96 | 2,74 | 43,22 | 80,30 |
|  | HD-ISS-1 | 20 | 45,54 | 35,00% | 0,14 | 56,85 | 12,95 | 3,95 | 42,90 | 88,20 |
|  | HD-ISS-2 | 46 | 43,38 | 41,30% | 0,20 | 49,13 | 12,93 | 12,30 | 43,48 | 87,89 |
|  | HD-ISS-3 | 16 | 47,37 | 56,25% | 0,18 | 45,50 | 11,31 | 15,44 | 43,00 | 93,84 |
| 24 MONTHS |  | 176 | F = 4.591,<br>p = 0.0015 | F = 1.404,<br>p = 0.2346 | F = 2.089,<br>p = 0.0843 | F = 12.57,<br>p < 0.0001 | F = 27,<br>p < 0.0001 | F = 87,<br>p < 0.0001 | F = 0.082,<br>p = 0.97 | F = 6.727,<br>p = 0.0004 |
|  | HC | 84 | 50,40 | 59,52% | 0,17 | 57,93 | 12,96 | 1,40 |  |  |
|  | HD-ISS-0 | 17 | 41,20 | 64,71% | 0,13 | 62,12 | 13,00 | 4,06 | 43,24 | 81,93 |
|  | HD-ISS-1 | 17 | 45,38 | 41,18% | 0,15 | 54,24 | 13,00 | 3,59 | 43,12 | 89,66 |
|  | HD-ISS-2 | 39 | 45,58 | 48,72% | 0,20 | 49,95 | 12,90 | 11,59 | 43,08 | 89,74 |
|  | HD-ISS-3 | 19 | 49,39 | 36,84% | 0,18 | 40,89 | 11,58 | 16,16 | 42,89 | 95,99 |

### fMRI data acquisition and preprocessing

Data were collected at four imaging sites with similar acquisition parameters. The details of the data acquisition were provided in (Tabrizi et al., 2011, 2009). The data was acquired using a 3T Siemens Trio TIM, Philips Intera, and Philips Achieva MRI scanners depending on the site. Regardless of the scanner the resting state fMRI scans were collected using repetition time (TR) = 3000 ms, echo time (TE) = 30 ms, flip angle = 80, field of view = 64 x 60 x 48, voxel size = 3.131, 165 volumes, for a scan duration of 8.25 minutes.

fMRI data were preprocessed with the *fMRIprep* v20.2.7 pipeline (Esteban et al., 2019; Markiewicz et al., 2024) (RRID:SCR_016216) during a data harmonization effort described in (Pustina et al., 2024). The full description of processing steps is provided in Supplementary Materials. In brief, the fMRI data were corrected for in-scanner motion, corrected for susceptibility-induced distortions using the acquired fieldmaps, band-pass filtered, co-registered to the T1-weighted structural scan, and resampled into several standard spaces including MNI152NLin2009cAsym, which was our space of reference for this study. Of note, we also applied the option for automatic removal of motion artifacts using independent component analysis (ICA-AROMA, Pruim et al. 2015) followed by spatial smoothing with an isotropic, Gaussian kernel of 6mm FWHM (full-width half-maximum). Corresponding “non-aggresively” denoised runs were produced after such smoothing. On the other hand, the “aggressive” noise-regressors were collected and placed in the corresponding confounds file. Several confounding time-series were calculated based on the *preprocessed BOLD*: framewise displacement (FD), DVARS, and three region-wise global signals. FD was computed using two formulations following Power et al. (2014): “absolute sum of relative motions” and Jenkinson et al. (2002): “relative root mean square displacement between affines”.

We combined two parcellation templates to extract the fMRI signals from the timeseries: (1) the Schaefer-200 parcellation template for cortical regions (100 parcels per hemisphere) (Schaefer et al., 2018) and the Tian-16 template for subcortical regions (Tian et al., 2020). We call this combined template the Schaefer-Tian parcellation. The cortical areas were assigned to 7 canonical resting state networks (RNSs) defined by (Yeo et al., 2011): visual (VIS), somatomotor (SOM), default mode network (DMN), dorsal attention network (DAN), fronto-parietal network (FPN), salience (SAL), limbic (LIM). The subcortical areas were categorized as subcortex (SCX) and further divided into striatum (STR), globus pallidus (GP) and thalamus (THAL), where needed.

### Structural connectivity

To estimate generative effective connectivity (GEC), we used a structural connectivity template derived from HCP dataset based on probabilistic tractography of diffusion weighted scans (DWI). The details of data acquisition are given in (Cabral et al., 2022). In brief, the connectome was constructed from diffusion- and T2-weighted MRI data of 32 healthy adults (mean age = 31.5 ± 8.6 years; 14 females) from the HCP. Diffusion MRI (89 min) was acquired on a custom high-gradient scanner. Diffusion data were reconstructed using a generalized q-sampling imaging algorithm in DSI Studio, and white-matter masks from T2 segmentations were co-registered to the b0 image via SPM12. Within each mask, 200,000 streamlines were sampled and nonlinearly warped to MNI space using diffeomorphic registration. Individual tractograms were aggregated into a normative MNI-space connectome representing healthy young adults and released within Lead-DBS toolbox. Consistent with the functional data, dense structural connectivity matrices were parcellated using the Schaefer-Tian template.

### AHBA Gene expression dataset

We used the Allen Human Brain Atlas (AHBA) gene expression dataset (Gryglewski et al., 2018; Hawrylycz et al., 2015, 2012) and the abagen python package (Arnatkevičiūtė et al., 2019; Hawrylycz et al., 2012; Markello et al., 2021) to investigate whether the anomalies observed in fMRI metrics at each brain region are spatially correlated with the *HTT* gene expression levels. The *HTT* expression levels available in AHBA were obtained postmortem from several donors and then averaged within each brain region. Consistent with the structural and functional data, we parcellated gene expression dataset using the Schaefer-Tian template (200 cortical and 16 subcortical parcels). The automatically generated text containing the details of the AHBA data processing is provided in supplementary materials.

### Graph theoretical analyses

Since the graph topology and underlying connectivity is expected to be different between GEC and FC, we computed and analyzed both GEC and FC matrices. We estimated several graph theoretical metrics that include modularity, clustering coefficient, density, efficiency, characteristic path length and assortativity. Modularity and clustering coefficient characterize whether the nodes tend to form clusters with each other. Density, efficiency, and characteristic path length quantify how efficient/fast information can transfer across distant nodes through the graph. Assortativity shows whether the nodes with similar characteristics tend to connect each other. We did not use rich club coefficient since we sought to characterize whole brain measures and hubs were not of interest in this study. Overall, these graph metrics characterize the segregation, integration and hierarchical organization of the connectivity, respectively. All graph metrics were computed using brain-connectivity toolbox (Rubinov and Sporns, 2010).

### Statistical analyses

Unless otherwise stated, we used linear mixed model (LMM) to make statistical inferences. We used multiple models to identify the best fit. The first, most basic model was:

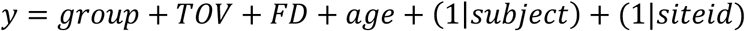

where group is the diagnostic variable (indicating healthy controls, HD-ISS-0, HD-ISS-1, HD-ISS-2 and HD-ISS-3), TOV indicates the time of visit (baseline, 12-months and 24-months), while we controlled for age and framewise-displacement (FD). To model longitudinal data, we included subject intercept as a random effect in the model. Finally, we included site ID intercept as another random effect to take into account shared variance due to different recording sites.

The alternative model we considered also included the interaction between group and study TOV variables to test longitudinal effects:

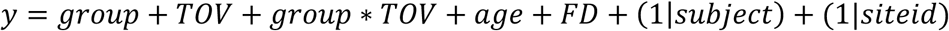

We compared the basic and alternative models with each other using likelihood ratio test where applicable. Unless the alternative models explained the variance significantly better than the simpler one, we used the simplest model without interaction term and random slope. Finally, we summarized the statistics for the estimates of interest using F-statistic and its associated p-value.

Regarding age, CAG, age x CAG interaction, total functional capacity (TFC), total motor score (TMS), symbol digit modality test (SDMT), we constructed the model:

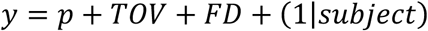

Where p is age, CAG, age x CAG, TFC, TMS or SDMT.

To correct for multiple comparison of connectivity strength analyses, we fit the model after shuffling HD-ISS and FD of the subjects and recorded the maximum F-statistic (ANOVA over the linear model) over 216 areas for 1000 permutations each (Winkler et al., 2016). For the rest of the multiple comparisons, we corrected using false discovery rate (FDR) approach (Benjamini and Yekutieli, 2001). The analyses were done using MATLAB.

### Network Based Statistics

To assess the differences in FC and GEC during disease progression, we performed network-based statistics (NBS). In brief, NBS approach addresses the multiple comparison problem for connectivity edges by correcting them at the network level (Zalesky et al., 2010). For EC, we used the directed version of the NBS toolbox. NBS approach came with computational complexity as it involves thousands of comparisons and their assessments with 5000 permutations. As shown later in the Results section, the longitudinal analyses did not yield useful results, and we focused on cross-sectional differences based on FCs obtained by averaging all sessions of each individual (i.e. collapsing time-of-visit). The same approach was used in previous studies using a subset of the TRACK-HD dataset (Odish et al., 2015). To investigate the effect of HD progression without the effect of normal aging, we modelled the aging effect in healthy controls using a linear model and regressed out this effect from the PwHD. Finally, rather than performing a separate ANOVA followed by post-hoc analyses between HCs and each HD-ISS stage, we directly performed T-tests between HCs and each HD-ISS group to assess the differences in network connectivity (i.e. HC vs. HD-ISS-0, HC vs. HD-ISS-1, HC vs. HD-ISS-2 and HC vs. HD-ISS-3). Since we made multiple comparisons (4 tests for each connection), we used a more conservative p-threshold of 0.01 rather than 0.05.

For networks that showed abnormalities in HD, we performed post-hoc tests to assess how each connection was related to known biomarkers of the disease, i.e., age, CAG length, symbol digit modality test (SDMT), total motor score (TMS) and total functional capacity (TFC). Similarly, we used LMMs to study these effects.

### Generative effective connectivity model

The model involves 216 nodes, where each node is coupled with each other via DWI-derived structural connectivity (SC) matrix. The local dynamics of each individual node are represented by normal form of a supercritical Hopf bifurcation. The details of the model are described elsewhere in the literature (Deco et al., 2023, 2017; Kringelbach et al., 2023)(see also Supplementary Materials).

To find the optimal GEC, we start with an empirical structural connectivity matrix, which is used for identifying directly connected areas ignoring the actual anatomical connectivity strength. At each iteration, we assess the similarity between empirical and model FCs, while adjusting the underlying EC. Then we use a heuristic approach to find the optimal GEC matrix that maximizes the similarity between empirical and model FCs. Therefore, instead of using FC that reflects the involvement of many complex dynamic processes such as interactions across local oscillatory activity, noise, global signal and various artifacts, we seek to infer an anatomically plausible connectivity structure that optimally explains the observed FC.

The two types of connectivity, the model-generated functional connectivity (*FC^model^*) and time-shifted functional connectivity (*FC^tau,model^*), were used to infer the optimal generative effective connectivity (GEC) matrix that explains both the observed functional connectivity (*FC^emprical^*) and time-shifted functional connectivity (*FC^tau,emprical^*). Tau for time-shifted FC is set to 1 TR (3 seconds). The details of the optimization procedure were provided in supplementary materials. During the optimization procedure none of the individuals reached the maximum number of iterations and all successfully converged to an optimal solution. This indicates that the optimization procedure was successful.

## Results

We investigated how potential resting-state fMRI biomarkers of HD can be characterized across various spatial scales within HD-ISS categories. First, we assessed the whole-brain graph theoretical measures of the FC and GEC networks. Then, we focused on FC and GEC strengths, characterizing the changes in global brain connectivity of each brain area. Finally, we performed an edge-level analysis based on the Network-Based Statistics (NBS) approach and further explored the relationships of the altered network with age, CAG, cognitive, and behavioral biomarkers. To understand the possible mechanisms behind the results, we studied the association between the huntingtin (*HTT*) gene expression map and FC strength.

### Data and Demographic Characteristics

Demographic characteristics for each group are shown in Table 1. The groups were balanced in sex, but not in age, i.e., HCs and HD-ISS-3 individuals were significantly older than HD-ISS-0, HD-ISS-1 and HD-ISS-2. Importantly, there were no significant differences between groups regarding framewise displacement (FD) (see Table 1). We observed a significant effect in FD across recording sites, with data from some sites having more in-scanner motion than in other sites, but the HD-ISS groups were generally well balanced at each site (Supplementary Materials). These observations influenced the choices we made for statistical analyses. Specifically, we used linear mixed models (LMM) given that most of the individuals had data from all three sessions. Approximately 33% of individuals had at least one session missing.

### Estimation of Generative Effective Connectivity

As described above, GEC has been shown to reveal rudimentary connectivity patterns that optimally generates the observed FC (Kringelbach et al., 2023). The model generates simulated FC of each participant by using a Hopf model, which relies on brain areas operating near the edge of noise-driven and oscillatory dynamics, also known as supercritical Hopf bifurcation. As an initial guess the model uses the average structural connectivity (SC) and connectivity weights are updated throughout the optimization only if two areas are structurally connected. We used a heuristic approach to find the GEC for each participant that minimizes the difference between observed FC and time-shifted FC and their simulated counterparts. For each participant the model successfully converged to an optimal solution with an average model fit between simulated and observed FC (Pearson-r = 0.7071 ± 0.0852).

### Graph theoretical measures

We constructed a linear mixed model to study the relationship between graph topology and HD-ISS-stage and time-of-visit (TOV), while controlling for framewise displacement. We found no significant differences in any of these measures between HD-ISS groups, time-of-visit (TOV), or their interaction (See Table 2). All graph theoretical measures except assortativity and density were significantly associated to framewise displacement. To further investigate this effect, we repeated the analyses after reducing the dimensionality of the data using PCA. We found that the first principal component (PC-1) was highly associated with framewise displacement and explained 88% of the variance in FC and 61% of the variance in EC, suggesting a substantial effect of in-scanner motion on the graph theory measures (See Supplementary Materials).

**Table 2.** Whole-brain graph metrics for FC and EC (assortativity, clustering coefficient, density, global efficiency, path length, modularity, and for EC). We summarized the F statistics as ANOVA of the proposed Linear Mixed Model. P-values are corrected for multiple comparisons using FDR. HD-ISS is the group variable (HC, HD-ISS-0..3), TOV indicates time-of-visit, age and framewise displacement are confounding variables. TOV x HD-ISS interaction is the alternative model to test whether longitudinal effects interact with the diagnosis. LR = Likelihood ratio of the alternative model explaining significantly greater variance than the reduced model.

|  | *Variable* | *HD-ISS* | *TOV* | *Age* | *Framewise Displacement* | *TOVxHD-ISS* |
| --- | --- | --- | --- | --- | --- | --- |
| *FC* | Assortativity | F=0.26,<br>p=1.000 | F=0.89,<br>p=0.849 | F=5.75,<br>p=0.247 | F=5.31, p=0.053 | F=1.91, p=1.000; LR=7.56, p=0.109 |
|  | Clustering coefficient | F=0.85,<br>p=1.000 | F=3.98,<br>p=0.228 | F=1.18,<br>p=0.682 | F=29.55, p<0.001 | F=0.94, p=1.000; LR=3.73, p=0.444 |
|  | Density | F=0.64,<br>p=1.000 | F=2.44,<br>p=0.437 | F=4.47,<br>p=0.257 | F=15.21, p<0.001 | F=0.47, p=1.000; LR=1.88, p=0.758 |
|  | Global efficiency | F=0.99,<br>p=1.000 | F=4.33,<br>p=0.228 | F=1.44,<br>p=0.680 | F=27.32, p<0.001 | F=1.05, p=1.000; LR=4.19, p=0.381 |
|  | Path length | F=0.92,<br>p=1.000 | F=4.22,<br>p=0.228 | F=1.57,<br>p=0.680 | F=29.20, p<0.001 | F=0.83, p=1.000; LR=3.32, p=0.505 |
|  | Modularity | F=0.43,<br>p=1.000 | F=1.37,<br>p=0.712 | F=1.65,<br>p=0.680 | F=26.83, p<0.001 | F=0.30, p=1.000; LR=1.19, p=0.879 |
| *GEC* | Assortativity | F=0.31,<br>p=1.000 | F=3.24,<br>p=0.532 | F=0.17,<br>p=1.000 | F=14.14, p<0.001 | F=0.93, p=1.000; LR=3.71, p=0.446 |
|  | Clustering coefficient | F=0.33,<br>p=1.000 | F=0.05,<br>p=1.000 | F=0.03,<br>p=1.000 | F=33.09, p<0.001 | F=0.95, p=1.000; LR=3.78, p=0.437 |
|  | Density | F=1.48,<br>p=1.000 | F=10.05,<br>p=0.024 | F=3.09,<br>p=1.000 | F=4.05, p=0.110 | F=0.72, p=1.000; LR=2.86, p=0.581 |
|  | Global efficiency | F=0.35,<br>p=1.000 | F=0.46,<br>p=1.000 | F=0.02,<br>p=1.000 | F=41.50, p<0.001 | F=0.75, p=1.000; LR=2.98, p=0.561 |
|  | Path length | F=0.51,<br>p=1.000 | F=1.81,<br>p=0.827 | F=0.36,<br>p=1.000 | F=32.30, p<0.001 | F=0.90, p=1.000; LR=3.59, p=0.465 |
|  | Modularity | F=0.82,<br>p=1.000 | F=1.48,<br>p=0.827 | F=0.98,<br>p=1.000 | F=35.80, p<0.001 | F=0.19, p=1.000; LR=0.74, p=0.946 |

### FC and GEC strengths

To characterize connectivity changes at the mesoscopic level, we studied the FC and GEC strengths of each cortical and subcortical area. As we did for graph theoretical measures, we constructed a linear mixed model to predict strength of each node by HD-ISS-stage, time-of-visit (TOV) and controlled for framewise displacement. We didn’t find any evidence suggesting that an alternative model involving HD-ISS x TOV interaction is significantly better for any brain area (See Supplementary Materials). After multiple comparison correction, we found no significant effects regarding time-of-visit, but we found several areas were significantly related to FD cross-sectionally (See Supplementary Materials).

One area showed significant differences between groups: left caudate (F-stat = 8.5, p-permutation < 0.001) in FC and in GEC (F-stat = 6.49, p-permutation = 0.012) (Supplementary Table) (Figure 1). These results suggested that caudate FC and GEC strength progressively decrease along HD-ISS.

**Figure 1.**
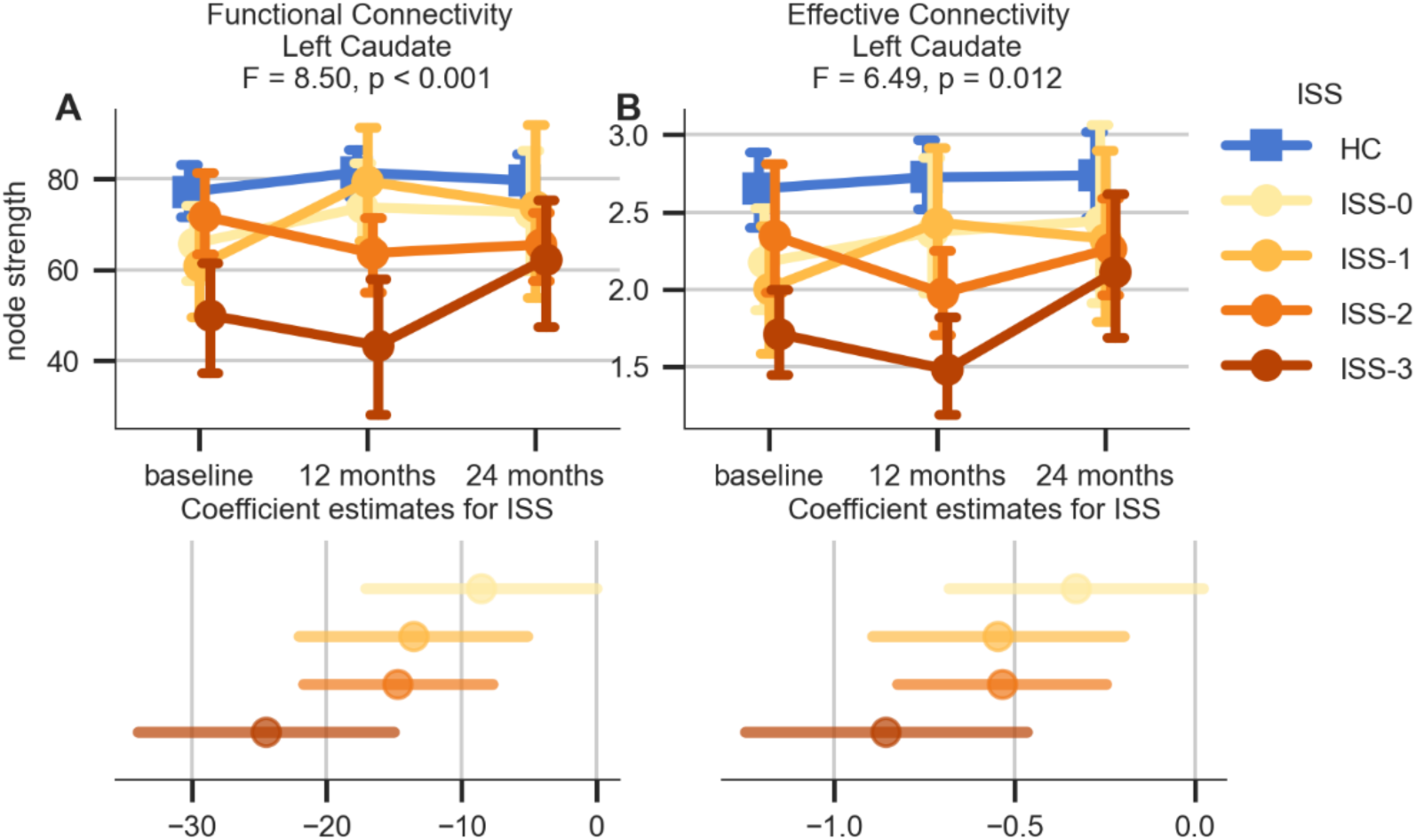
FC and GEC strength (global brain connectivity) alterations for FC left caudate (A), GEC left caudate (B). Upper panels show boxplots for each group across time of visit. Lower panels show the coefficient estimates of the linear mixed model for each HD-ISS category taking HC as reference. Tails show the 95% confidence intervals.

### Network-Based Statistics

We performed pairwise T-tests between FCs and ECs of healthy controls and each HD-ISS group at edge level and controlled for framewise displacement as a confounding variable. We found no differences between HCs versus HD-ISS-0 and HD-ISS-1 individuals. We found that the connections were decreased in HD-ISS-2 compared to controls, and these connections were all among caudate nuclei and other subcortical nuclei (Table 3). In HD-ISS-3, we found more widespread decreased connectivity and the majority of the decreased connections were still linked to the striatum. When we performed NBS on GEC matrices, we found similar patterns in HD-ISS-3 that were linked to impaired connections of the caudate (Figure 3), although this time the most impaired caudate connectivity was towards anterior brain areas. The percentage of decreased connections to or from the striatum were higher when measured with GEC than when measured with FC (Figure 2). Overall, these results indicate that the impairments in large-scale connectivity are noticeable only after HD-ISS-2 and they are mostly in cortico-striatal connections.

**Figure 2.**
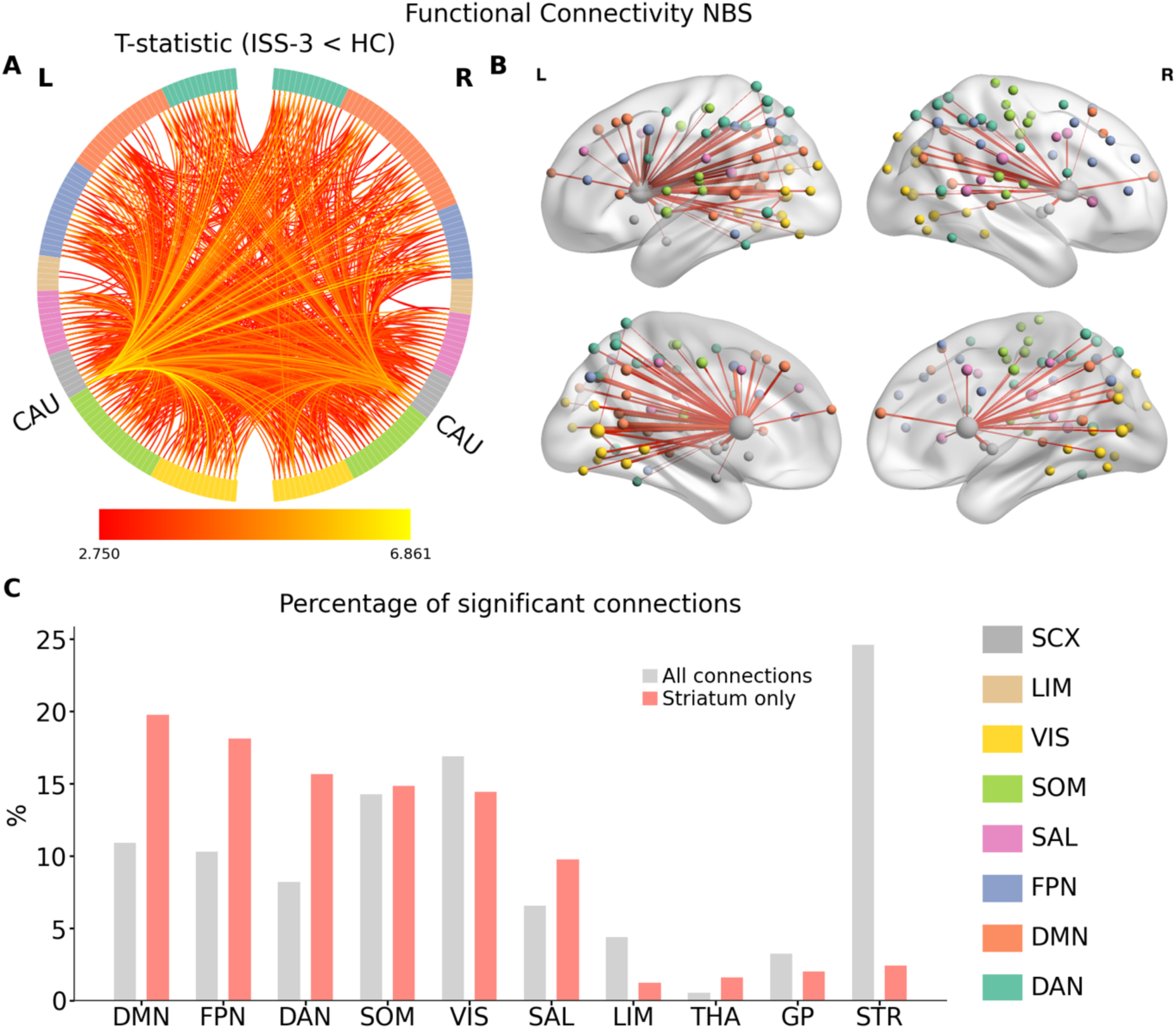
FC alterations based on NBS approach. Upper panel: Significantly different network edges (p < 0.01) (HD-ISS-3 < HC) illustrated in circle plot (left) and brain plot (right). Lower panel: The percentage of significant connections for each resting state network, thalamus, globus pallidus and striatum (caudate and putamen). Gray bars indicate total percentage of significant connections, whereas orange bars indicate only caudate and putamen.

**Figure 3.**
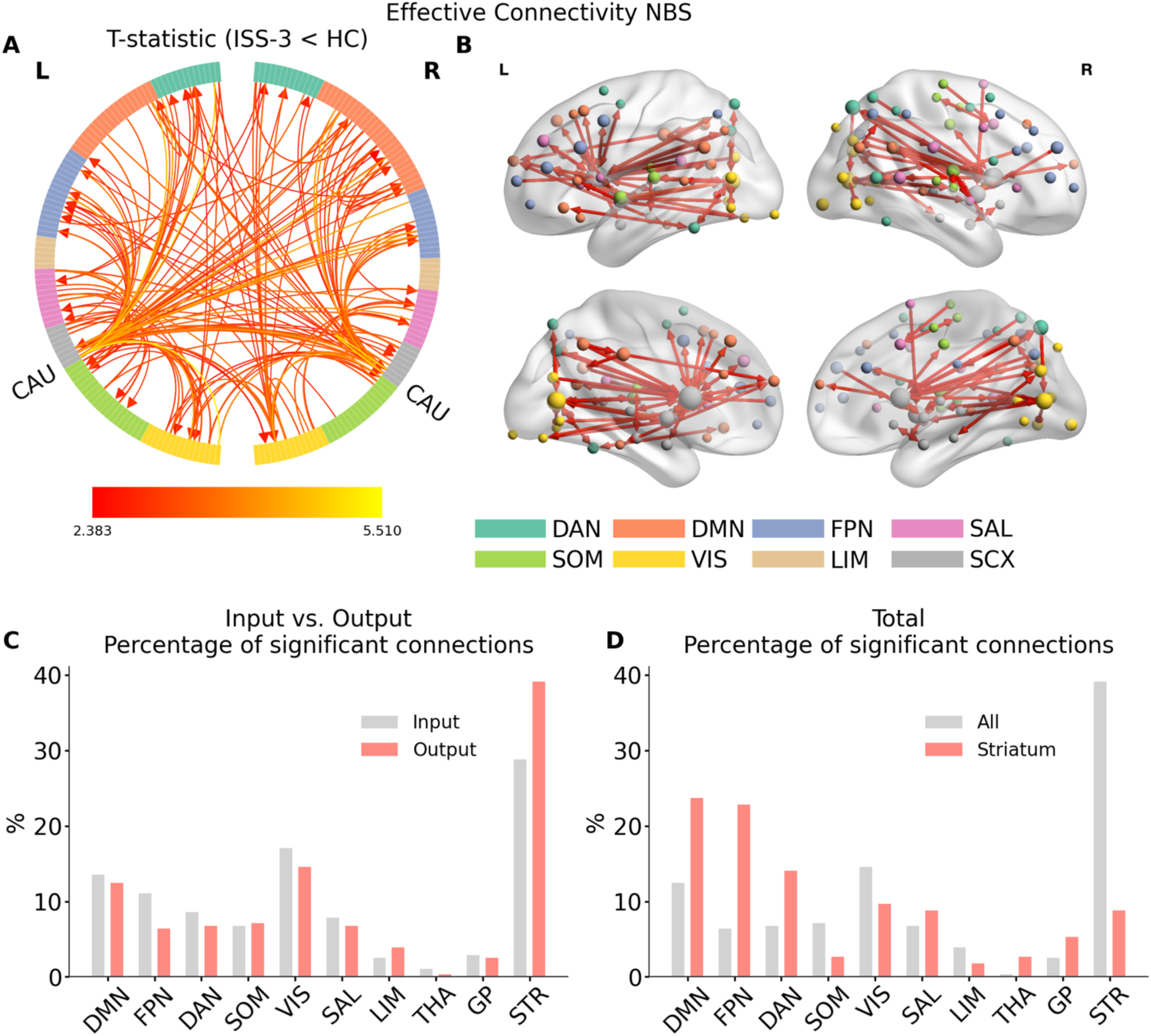
Effective Connectivity alterations based on NBS approach. Upper panel: Significantly different network edges (p < 0.01) (HD-ISS-3 < HC) illustrated in circle plot (left) and brain plot (right). Lower panel: The percentages of input (gray) and output (orange) connections to/from each resting state network and subcortical hub (Left). The percentages of significant connections for each resting state network and subcortical hub (right). Gray bars indicate total percentage of significant connections, whereas orange bars indicate only caudate and putamen.

**Table 3.** Significantly altered networks in HD-ISS-2 category.

| Functional Connectivity |  |  | Effective Connectivity |  |  |
| --- | --- | --- | --- | --- | --- |
| Edge | T-statistic | P-value (FDR corrected) | Edge | T-statistic | P-value (FDR corrected) |
| CAU_R - CAU_L | 5,564175 | 2,13E-07 | CAU_L - PUT_L | 5,580143 | 1,28E-07 |
| PUT_L - CAU_L | 5,518817 | 2,13E-07 | PUT_L - CAU_L | 5,226926 | 3,22E-07 |
| CAU_R - GP_L | 5,33058 | 3,37E-07 |  |  |  |
| PUT_R - CAU_R | 5,23805 | 3,83E-07 |  |  |  |
| GP_L - CAU_L | 4,557019 | 5,78E-06 |  |  |  |

### Role of age and CAGs

We used a linear regression model to investigate whether the connections with decreased strength in HD-ISS-3 were related to age, CAG, or the age × CAG interaction. Results were corrected for multiple comparisons using the false discovery rate. Age-related functional decreases in connectivity were characterized by connections between caudate and DMN nodes as well as sensory networks (visual and somatomotor). CAG was related largely to altered connections to and from the striatum (caudate and putamen). The age × CAG interaction was related to decreased connections in FPN and globus pallidus (Figure 4).

**Figure 4.**
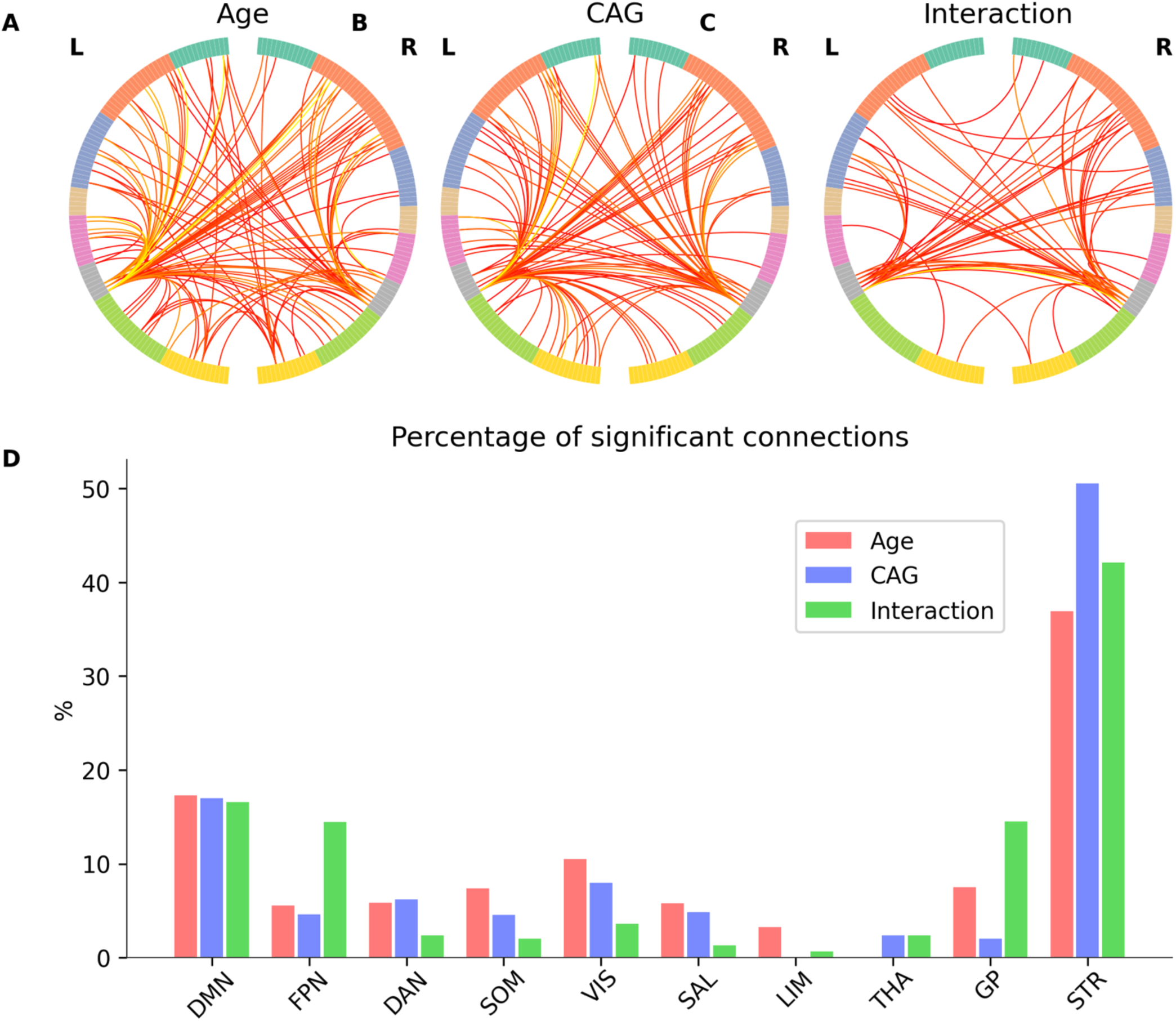
The association between the connections within the altered network (significant network by NBS), age, CAG and age x CAG interaction. Upper panel: Connections with significant coefficient estimate t-statistic for age (A), CAG (B) and Age x CAG interaction (C). Lower panel: The percentage of significant connections for each resting state network, thalamus, globus pallidus and striatum (caudate and putamen) for age (red), CAG (blue) and age x CAG interaction (green).

### Relationship to cognitive measures

We also studied the relationship between FC alterations and cognitive measures. Similar to age and CAG, we used linear mixed model to study the association between SDMT, TMS and TFC in the significantly altered network after controlling for framewise displacement. Both SDMT and TFC had widespread associations with FC connections (Figure 5). In contrast, TMS exhibited a sparse pattern such that only a few connections between caudate and specific cortical areas correlated with it.

**Figure 5.**
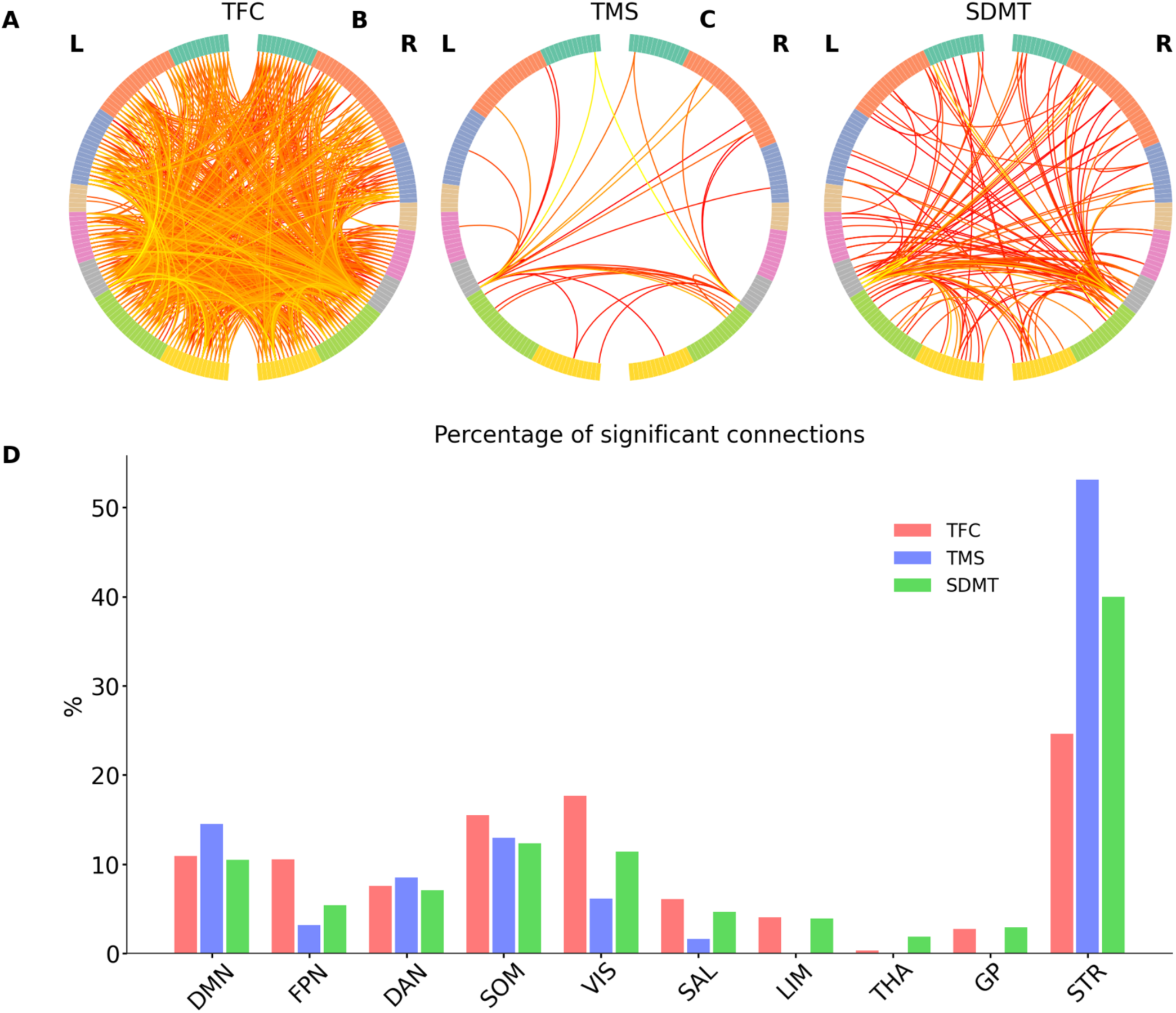
The association between the connections within the altered network connections in HD-ISS-3 and cognitive indicators: total functional capacity (TFC), total motor score (TMS) and SDMT. Upper panel: Connections with significant coefficient estimate t-statistic for TFC (A), TMS (B) and SDMT (C). Lower panel: The percentage of significant connections for each resting state network, thalamus, globus pallidus and striatum (caudate and putamen) for TFC (red), TMS (blue) and SDMT (green).

### Association between observed alterations and gene expression topography of HTT

After identifying the alterations in brain connectivity, we checked whether the spatial pattern of these anomalies follows the topography of *HTT* gene expression patterns in the brain from the AHBA database. Following a similar approach as in Preller et al. (Preller et al., 2018), we calculated the Spearman correlation between the *HTT* gene expression map and the map of FC anomalies across the brain. To understand how this relationship differed across HD-ISS categories, we used the difference of T-statistics of the coefficient estimates for each HD-ISS group with respect to those of healthy controls (HCs). We found a significant association with *HTT* map only in HD-ISS-2 and HD-ISS-3 (Figure 6), but not in HD-ISS-0 and HD-ISS-1.

**Figure 6.**
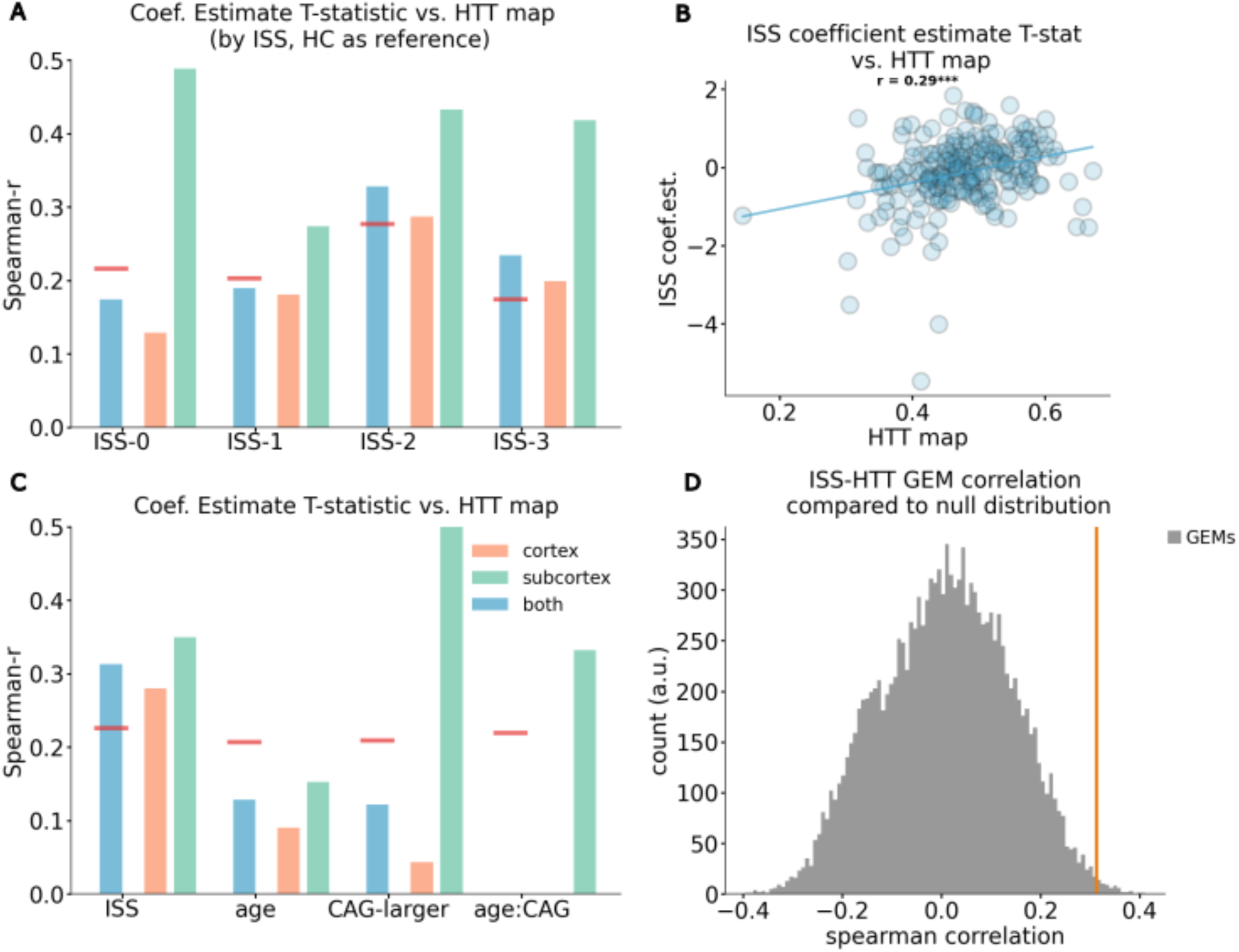
The relationship between the changes in FC strength and *HTT* gene expression map. A. The spearman correlation between *HTT* map and the coefficient estimate T-statistic for each HD-ISS category (deviation from HC as reference). B. Scatter plot showing the relationship between HD-ISS coefficient estimate T-statistic and *HTT* map. C. The spearman correlations between *HTT* maps and the coefficient estimate T-statistic for HD-ISS, age, CAG and age x CAG interaction. Blue bars indicate whole-brain correlations, salmon bars indicate cortex-only and green bars indicate subcortex-only correlations. Red lines indicate the 95th percentile along the distribution of correlation coefficient for the other 15k gene expression maps (D).

## Discussion

The primary goal of our study was to identify resting-state fMRI biomarkers using advanced models of brain connectivity. We pursued this goal by analyzing longitudinal data from 231 participants in the Track-On HD study. We did not find significant anomalies related to longitudinal changes across different HD-ISS progression groups. We then combined the longitudinal data using a method known to enhance analysis robustness (Cho et al., 2021; Odish et al., 2015) and found results consistent with known HD pathology, namely decreased connectivity in key regions associated with HD neuropathology, such as the caudate and putamen. These anomalies were detected only in later stages of HD (HD-ISS-2, HD-ISS-3), making them less useful as biomarkers in earlier stages of progression (HD-ISS-0, HD-ISS-1). We will revisit some of these findings in the following paragraph and compare them to previous reports with similar or conflicting findings.

Connectivity strength, beyond being a graph metric, has also been useful in quantifying overall brain connectivity, specifically how each area connects to the rest of the brain (Anticevic et al., 2014; Cole et al., 2010). We did not observe any differences in the whole-brain graph theoretical measures of the FC and GEC networks, suggesting that HD does not lead to widespread restructuring of functional dynamics across the entire brain. Instead, we found that FC and GEC strengths were only altered in the striatum. The edge-level analysis showed no significant differences between HD-ISS-0 or HD-ISS-1 and healthy controls, but such differences appeared in later stages. In HD-ISS-2, connectivity impairments were confined within subcortical striatal regions, whereas in HD-ISS-3 we observed more widespread reduced connectivity between the caudate, globus pallidus, and the rest of the cortex. Additionally, there was a significant linear decrease in bilateral connectivity in the caudate across the HD-ISS progression groups, consistent with a gradual loss of function during HD progression. The spatial correlations between *HTT* expression levels and FC anomalies also appeared in HD-ISS-2 and HD-ISS-3, but not in earlier stages. Overall, these findings indicate that FC anomalies become easier to detect as HD progresses and clinical signs emerge.

We investigated how the connectivity differences between caudate and cortex were associated with other markers of HD (i.e. age and CAG lengths) and clinical outcome measures (SDMT, TMF and TFC). The results suggested that age-related connectivity impairments were higher in cortex, especially in sensory networks and globus pallidus, whereas CAG-length-related impairments affected the striatum more than the cortex. Thus, age and CAG-related pathologies have distinct but related effects on the brain. CAG repeats might be leading the deterioration of the connectivity in striatum because of the related neurodegeneration, while impairments in DMN and sensory network connectivity might be a more generic signature of aging as these networks are known to be affected in healthy aging.

While our findings are intuitive and often in line with the known HD pathology, they are not fully aligned with the previous literature. For example, previous studies reported altered graph topology in HD in a variety of metrics (Gargouri et al., 2016; Harrington et al., 2015; McColgan et al., 2017). Harrington et al. (2015) reported that the efficiency and rich club coefficients is reduced in HD, whereas no significant differences were observed in other metrics including connectivity strengths. Similarly, Gargouri et al., (2016) used a subset of the Track-On HD dataset and found significant reductions in characteristic path length. We sought to compare how these measures are aligned along HD-ISS categorization but did not find any significant differences in graph topology, which might be due to differences in fMRI data processing or, more simply, the adoption of the new HD-ISS progression system. Since HD-ISS stages provides a different split of participants within the traditional pre-HD group, this might alter the outcome both by the slightly different phenotypic categorization and by the inadequate sample size in some categories. On a similar note, the review by Pini and colleagues (Pini et al., 2020) concluded that HD affects FC in visual, sensory-motor, executive and frontoparietal areas. In sensory-motor areas, Pini et al. noticed a non-linear relationship with disease progression: FC is first decreased below healthy control levels upon CMD and then abnormally increased as the disease progresses. Our results did not identify these cortical patterns and were almost entirely localized around the caudate, globus pallidus and putamen. It is worth noting that our results stem from the investigation of a single dataset while Pini et al. reviewed multiple studies before reaching their conclusions.

We also believe that the effect of in-scanner motion has a substantial impact on the discrepancies between previous findings since we found that ∼80% of the variance in graph measures is related to in-scanner motion. It is possible that motion effects are propagated downstream to the results in different ways, depending on how motion is corrected and accounted for by each study, as well as the amount of motion at each site or study. Indeed, we found systematic differences of in-scanner motion between study sites, which suggests that data obtained from one site/study may not be equivalent to data obtained from another site due to different conditions that lead to different motion levels. Regarding the correction methods, we applied standard coregistration of volumes and kept framewise displacement as nuisance variable in each analysis. We also controlled for study site in our analyses. On the other hand we avoided more aggressive noise removal strategies such as global signal regression to avoid the introduction of artifactual anti-correlations in model-generated FC (Murphy et al., 2009). From a statistical perspective, we used a stringent approach to compensate for the inflation of type-I errors due to multiple comparisons. It is worth noting that despite these discrepancies, previous studies that utilized the same Track-On HD dataset showed similar findings to ours, i.e., weak or null results on functional connectivity measures (Dumas et al., 2013; Estevez-Fraga et al., 2025; Odish et al., 2015). Thus, our results are consistent with other results reported on the same dataset despite the differences on data processing and participant classification.

In our findings, TFC and SDMT exhibited a widespread relationship with altered networks, whereas TMS was more tightly related to striatal disconnections. This pattern of findings is consistent with the nature of these measurements. TFC is a general measure of everyday function and is likely to be related to the integrity of multiple brain networks, any of which can impair an activity in everyday functions. SDMT is also a cognitive measure that includes a motor component to it, potentially requiring multiple brain networks of attention, memory, and motor planning. TMS on the other hand is strictly focused on motor ability, and for that reason can be more narrowly related to the motor planning networks that include the striatum.

This study has both strengths and limitations. We performed data processing and analyses that are cutting-edge and followed proper standards of statistical thresholding. We also integrated and cross-analyzed three different domains: imaging, clinical outcome measures, and gene expression data. The data we used come from Track-On HD, a study known for high-quality data across all domains (Pustina et al., 2024). However, our attempt to apply the HD-ISS framework to this dataset resulted in unbalanced groups, which limited the statistical power. Systematic analyses of fMRI data indicate that optimal group sizes should be N=40 or more (Ma et al., 2024), while our smallest HD-ISS group had N=13. Another potential limitation might be insufficient duration of fMRI scans and the low sampling frequency. The Track-On HD fMRI data was collected for 8.25 minutes with a sampling frequency of 3 seconds. It is well known that longer fMRI sessions yield more reliable results (see review by Noble et al., 2019). For example, Birn et al. (Birn et al., 2013) observed gradual improvements in test-retest reliability when scans were extended from 5-7 minutes, with around 16 minutes being optimal. Recent recommendations for other disorders also suggest rs-fMRI sessions should last between 13 and 25 minutes (Caeyenberghs et al., 2024). Higher temporal sampling rates (< 3 seconds) can also increase the number of data points and improve the ability to detect group differences (Birn et al., 2013). Our findings suggest that future rs-fMRI studies in HD should include longer scans at higher sampling rates, such as 1 second.

In conclusion, we found that brain connectivity anomalies became increasingly detectable with more advanced HD-ISS stages. We also showed that these anomalies correlate with clinical scores and the spatial distribution of the *HTT* gene transcription. Work remains to be done to disentangle the methodological limitations mentioned above from the true effect of HD pathology in order to obtain sensitive biomarkers.

